# Human blood contains circulating cell-free mitochondria, but are they really functional?

**DOI:** 10.1101/2020.12.29.424785

**Authors:** Antoine Stier

## Abstract

Dache et al. (2020, *FASEB J*. **15**, e2002338–15) recently reported the presence of *respiratory-competent* cell-free mitochondria in human blood (up to 3.7 × 10^6^ per mL of blood), providing exciting perspectives on the potential role of these extra-cellular mitochondria. While their evidence for the presence of cell-free mitochondria in human blood is compelling, their conclusion that these cell-free mitochondria are respiratory-competent or functional has to be re-evaluated. To this end, we evaluated the functionality of cell-free mitochondria in human blood using high-resolution respirometry and mitochondria extracted from platelets of the same blood samples as positive controls. While cell-free mitochondria were present in human plasma (*i*.*e*. significant MitoTracker Green Fluorescence and complex IV activity), there was no evidence suggesting that their mitochondrial electron transport system (ETS) was functional (*i*.*e*. respiration rate not significantly different from 0; no significant responses to ADP, uncoupler or mitochondrial inhibitors oligomycin and antimycin A). Yet, *in vitro* complex IV activity was detectable and even slightly higher than levels found in mitochondria extracted from platelets, suggesting that cell-free mitochondria in human blood are likely to only retain a non-functional part of the electron transport system. Despite being unlikely to be fully functional in the narrow-sense (*i*.*e*. capable of oxidative phosphorylation), circulating cell-free mitochondria may have significant physiological roles that remain to be elucidated.

## Introduction

Mitochondria are the powerhouse of eukaryotic cells using oxidative phosphorylation to convert nutrients into ATP. They also have key roles in cell signalling (1) and ageing (2). Vertebrates possess functional mitochondria in every cell type, with the notable exception of mammalian mature erythrocytes (3). Dache et al. (2020) recently reported the presence of cell-free mitochondria in human blood, providing exciting perspectives on the role of these extra-cellular mitochondria, and their potential to be used as prognosis tool for various diseases (4). While their evidence for the presence of cell-free mitochondria in human blood is compelling and has been supported by other studies (*e*.*g*. (5, 6)), their conclusion that these cell-free mitochondria are respiratory-competent or functional has to be re-evaluated (7) since this is critical to uncover the physiological role they may or may not play (8).

Indeed, the interpretation that circulating cell-free mitochondria are functional is exclusively based on the authors’ graphical interpretation of *in vitro* O_2_ consumption rates (Fig. 7 in (4)), and not on statistical testing considering their sample size was N = 1 (despite having n = 3 replicates of the same pooled sample; Dache et al. *pers. com*.). Based on the data being presented in (4), it seems impossible to conclude that circulating cell-free mitochondria are functional, as recently highlighted by (7). The *in vitro* respirometry assay used in (4) to assess functionality has several limitations. First, the addition of ascorbate and TMPD to test complex IV maximal capacity indeed elicits a rise in O_2_ consumption, but no negative control is shown by the authors, while there is a known chemical O_2_ consumption linked to the auto-oxidation of ascorbate and TMPD (9). Second, there is no visible inhibition of O_2_ consumption by antimycin A (*i*.*e*. a complex III inhibitor), suggesting that the minor O_2_ consumption being observed is unlikely to be of mitochondrial origin. Third, the role of mitochondria is not to consume O_2_ *per se*, but to produce ATP, and no test of oxidative phosphorylation functionality (*e*.*g*. stimulation of respiration by ADP addition, or inhibition of respiration by oligomycin addition, an inhibitor of ATP synthase) is provided (7).

Two alternative explanations could explain the apparent lack of functionality of these circulating cell-free mitochondria. It is possible that circulating cell-free mitochondria are not functional because they are damaged, or that only a small fraction of them (*i*.*e*. potentially the ones that arrived in the blood stream only very recently) are still somewhat functional. Mitochondrial integrity might be evaluated to some extent by the addition of exogenous cytochrome c during respirometry assay. Cytochrome c does not penetrate intact mitochondria but penetrates within partially damaged ones eliciting a rise in O_2_ consumption (10). Alternatively, it is possible that circulating cell-free mitochondria are functional, but that the *in vitro* respirometry assay of (4) was not well suited for these mitochondria (7). Indeed, the mitochondrial uncoupler FCCP has been added directly to the respiration medium at a concentration of 4μM. However, uncouplers like FCCP have a narrow optimal working range and require a progressive titration (*e*.*g*. 0.5μM steps; (10)). Indeed, when added in excess, uncouplers can partially to fully inhibit mitochondrial respiration, and partially damaged mitochondria are likely to be more sensitive to such phenomenon than intact ones (10). Consequently, it is possible that the cell-free circulating mitochondria observed in (4) were functional, but that they were fully inhibited by an inappropriate amount of mitochondrial uncoupler in the respiratory medium.

A proper assessment of the functionality of circulating cell-free mitochondria is therefore required to establish if they are really functional (7), or potentially non-functional remnants of mitochondria that have been extruded by cells in the blood stream. In this study, we aim to provide elements to answer this question by characterizing mitochondrial bioenergetics properties of circulating cell-free mitochondria using high-resolution respirometry and mitochondria extracted from platelets of the same blood samples as positive controls.

## Methods

### Sample collection

This study was performed in accordance with the WMA Declaration of Helsinki. Blood samples were collected at the University of Turku by a trained and habilitated lab technician from six healthy volunteers (2 males and 4 females) following signature of an informed consent form. Approximately 80mL of blood was taken from each volunteer using BD vacutainer K_2_EDTA tubes, which were progressively cooled and stored on ice before processing (max. 1 hour).

### Sample processing

We used a protocol similar to the one used by (6) to isolate circulating cell-free mitochondria. Blood samples were first centrifugated at 150*g* for 20min (4°C) to isolate platelet-rich plasma (PRP) from other blood cells. PRP was subsequently centrifugated at 2500*g* for 30min (4°C) to pellet platelets and obtain cell-free plasma. Mitochondria were isolated from platelets pellet following (11), with 4 cycles of 10 passes in glass/glass Potter-Elvehjem pestle homogenizer and 2500*g* centrifugation (4°C, 1min) to re-form platelet pellet and facilitate cellular membrane disruption by homogenization. Platelet samples were then centrifugated at 800*g* (4°C, 10min) and the supernatant containing isolated mitochondria was subsequently processed along with cell-free plasma. Both sample types were centrifugated at 10000*g* (4°C, 10min) to pellet mitochondria. Mitochondrial pellets were resuspended in 2mL of Mir05 respiration buffer (0.5 mM EGTA, 3 mM MgCl_2_, 60 mM K-lactobionate, 20 Mm taurine, 10 mM KH_2_PO_4_, 20 mM Hepes, 110 mM sucrose, free fatty acid bovine serum albumin (1 g/L)), and subsequently washed by centrifugation at 10000*g* (4°C, 10min). Mitochondrial pellets were finally resuspended in 100μL of Mir05. Mitochondrial proteins were quantified using the Biuret method, accounting for the fact that Mir05 contains 1mg.mL^-1^ of BSA. Mitochondrial content was measured using MitoTracker Green fluorescent probe (Invitrogen). Mitochondria were incubated in Mir05 supplemented with 200 nM of MitoTracker Green for 30min at 37°C, washed by centrifugation at 10000*g* (4°C, 10min) and re-suspended in 250μL of Mir05 before being transferred in 96-well black microplates (duplicates of 100μL) for fluorescence signal acquisition using an Enspire 2300 multilabel plate reader. Positive (standard dilution of *Saccharomyces cerevisae*) and negative (Mir05 and unstained cells) controls were run on each plate. MitoTracker Green data is expressed as relative fluorescence per mg of mitochondrial protein (see Fig. 1).

**Fig. 1:**
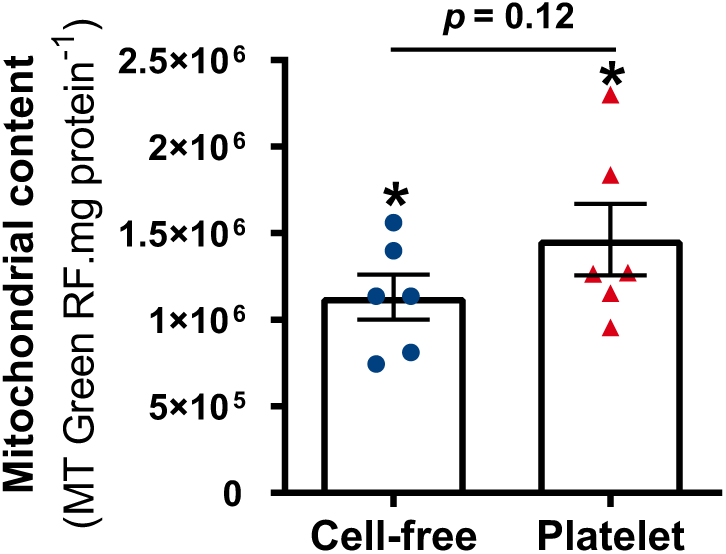
Mitochondrial content evaluated with MitoTracker Green relative fluorescence between circulating cell-free mitochondria and mitochondria extracted from platelets. Significant difference from 0 for each sample type is indicated by an *. Mean values are plotted ± 95% C.I., N = 6.

### High-resolution respirometry assay

Mitochondrial respiration was assessed using the high-resolution respirometry system O2k (Oroboros Instruments, Innsbruck, Austria) at 37°C using *ca*. 100 μg (range: [75; 125], cell-free mitochondria = 93.4 ± 7.5 μg, platelet mitochondria = 108.2 ± 4.9 μg, *p* = 0.12) of mitochondrial protein per assay (for comparison, 64μg of mitochondrial protein was used in (4)). Mitochondria were added to the respirometry chambers containing 2mL of Mir05 pre-equilibrated at 37°C and chambers were closed after 2min of equilibration. A substrate-uncoupler-inhibitor titration (SUIT) protocol was used to assess the functionality of different components of the mitochondrial electron transport system. Substrates of complex I (P: pyruvate 5mM and M: malate 2mM) were first added, followed by a saturating amount of ADP (2mM). Substrate of complex II (S: succinate 10mM) was then added to stimulate mitochondrial respiration fueled by both complexes I and II. Cytochrome c (Cyt c: 10μM) was added to test potential issues linked to mitochondrial inner membrane integrity. ATP synthesis was then inhibited with oligomycin (O: 2.5µM). A progressive titration (0.5μM steps) of the uncoupler FCCP was subsequently conducted until no further stimulation could be observed (maximum stimulation was reached at *ca*. 1.5μM), at which stage mitochondrial respiration was fully inhibited using the complex III inhibitor antimycin A (AA: 2.5µM). Finally, the maximal activity of complex IV was assessed using ascorbate (Asc: 2mM) and N,N,N’,N’-tetramethyl-p-phenylenediamine (TMPD: 0.5mM). Since there is a known chemical O_2_ consumption linked to the auto-oxidation of ascorbate and TMPD (9), 2 negative control samples (*i*.*e*. 50µL of Mir05 instead of mitochondria) were analyzed following the full SUIT protocol, so chemical O_2_ consumption could be subtracted from ascorbate+TMPD values of biological samples (see Fig. 2A).

**Fig. 2:**
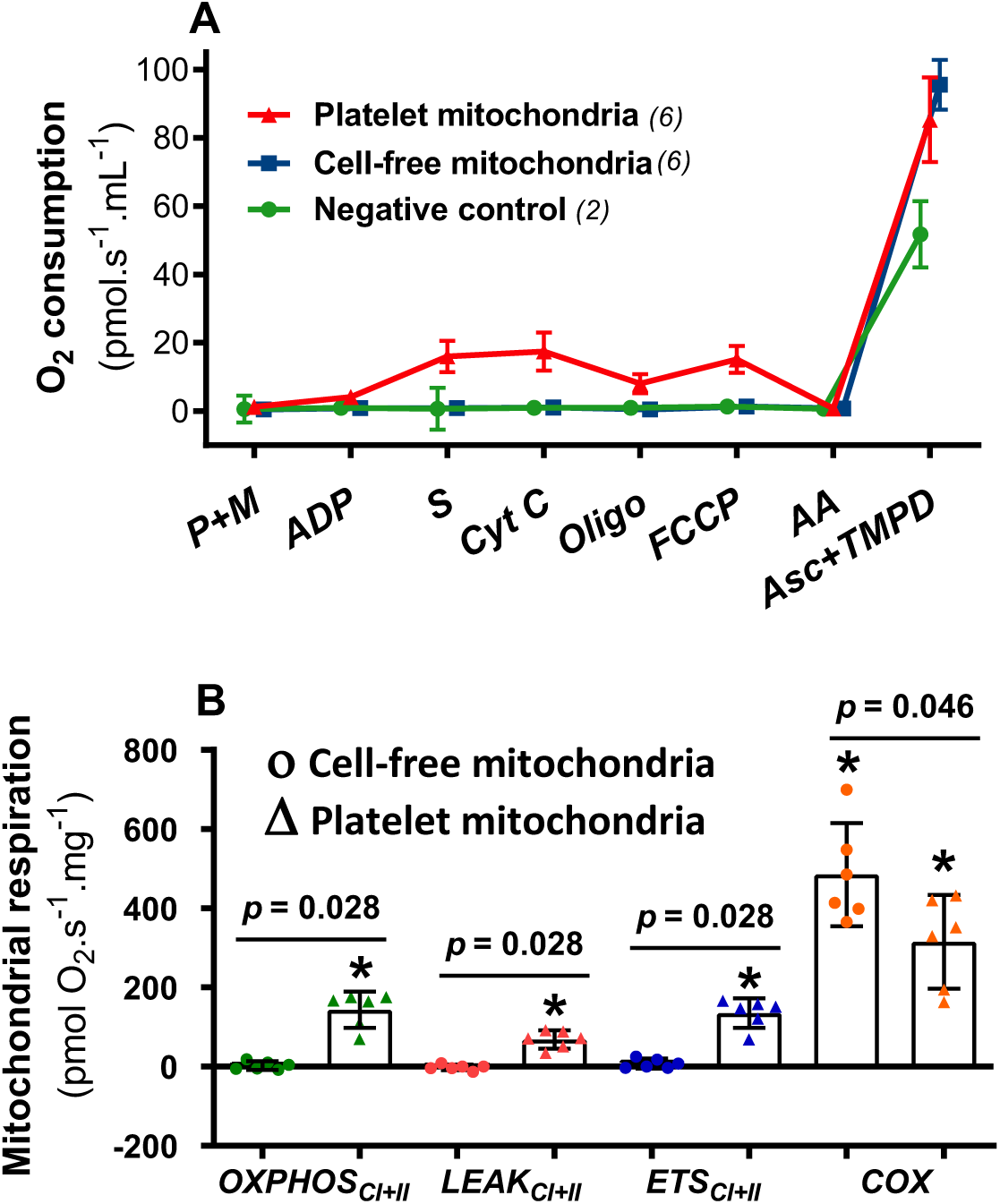
Mitochondrial bioenergetics properties of human circulating cell-free mitochondria and mitochondria isolated from platelets. **(A) Responses to a typical substrate-uncoupler-inhibitor protocol evaluated by high-resolution respirometry assay** (see methods for full details, and results for information on statistics). Negative control samples consisted of 50µL of respiration buffer instead of mitochondria. While platelets exhibited a typical response to SUIT protocol (all p < 0.05), cell-free mitochondria only responded to Asc+TMPD (p < 0.001, all other p = 1.00). **(B) Mitochondrial respiration rates of platelet vs. cell-free mitochondria**. Significant difference from 0 for each parameter is indicated by an *. All mean values are plotted ± 95% C.I., N = 6 in every case, except for negative controls (N = 2).

### Data analysis

Difference in mitochondrial content (MitoTracker Green relative fluorescence) between cell-free and platelet mitochondria was analyzed using non-parametric Wilcoxon paired test, and difference to 0 for each sample type using non-parametric one-way Wilcoxon test.

Oxygen consumption rates (pmol.s^-1^.mL^-1^) were extracted using DatLab 7.0 (Oroboros Instruments, Innsbruck, Austria) after correction for background noise. Responses to SUIT protocol of both plasma and platelet mitochondria were analyzed using a linear mixed model with sample type (cell-free *vs*. platelet), stage (P+M, ADP, S, Cyt C, O, FCCP, AA, Asc+TMPD) and their interaction as fixed factor; while controlling for pseudo-replication by including stage(ID) as a random factor to account for the fact that plasma and platelet samples came from the same individual (ID), as well as that measurements were repeated over the same samples in the different stages of the SUIT protocol. Since a significant interaction between sample type and stage was detected (see results), data was subsequently analyzed for each sample type individually to assess the differences between stages using Bonferroni-corrected post-hoc tests.

Mitochondrial respirations rates were calculated by subtracting non-mitochondrial oxygen consumption (after AA inhibition) to specific stages (*OXPHOS*_*CI+II*_ = S - AA; *LEAK* _*CI+II*_ = O - AA; *ETS* _*CI+II*_ = FCCP - AA), and by subtracting chemical O_2_consumption (*i*.*e*. obtained from negative controls, see Fig 2A) to Asc+TMPD values (*COX* = Asc+TMPD - AA - chemical O_2_ consumption). Those values were corrected by the quantity of mitochondrial protein used in each assay and therefore expressed as pmol.s^-1^.mg^-1^. Differences between cell-free and platelet mitochondria for each parameter were analyzed using non-parametric Wilcoxon paired tests, and differences to 0 of each parameter were tested using non-parametric one-way Wilcoxon tests. All analyses were conducted using SPSS 25.0, p-values < 0.05 were considered significant and values are expressed as mean ± 95% C.I.

## Results

There was a significant MitoTracker Green signal in both circulating cell-free (*p* = 0.023) and platelet (*p* = 0.023) mitochondria samples (Fig. 1). Mitochondrial content was lower (by *ca*. 33%) in circulating cell-free than in platelet mitochondria, but not significantly so (*p* = 0.12, Fig. 1).

Responses to SUIT protocol of both cell-free and platelet mitochondria, as well as negative controls, are presented in Fig. 2A. There was a significant interaction between sample type and stage in the way mitochondria responded to the SUIT protocol (*F*_*7,40*_ = 16.99, *p* < 0.001). Platelet mitochondria responded in the expected way to the SUIT protocol (*F*_*7,8*.*1*_ = 84.4, *p* < 0.001; Fig. 2A), with an increase in O_2_ consumption following ADP addition (P+M *vs*. ADP: *p* = 0.012), stimulation by complex II substrate S (ADP *vs*. S: *p* = 0.005), non-significant effect of Cyt c addition (S *vs*. Cyt c: *p* = 1.00), a decrease in O_2_ consumption following ATP synthase inhibition (Cyt c *vs*. O: *p* = 0.042), stimulation by FCCP titration (O *vs*. FCCP: *p* = 0.028), full inhibition by complex III inhibitor AA (FCCP *vs*. AA: *p* = 0.007) and clear stimulation by fueling directly complex IV with electrons (AA *vs*. Asc+TMPD: *p* < 0.001). On the contrary, although cell-free mitochondrial O_2_ consumption was affected by SUIT stage (*F*_*7,7*.*9*_ = 159.1, *p* < 0.001; Fig. 2A), they did not significantly respond to the SUIT protocol (P+M *vs*. ADP: *p* = 1.00; ADP *vs*. S: *p* = 1.00; S *vs*. Cyt c: *p* = 1.00; Cyt c *vs*. O: *p* = 1.00; O *vs*. FCCP: p = 1.00; FCCP *vs*. AA: *p* = 1.00), except for the clear stimulation obtained by fueling directly complex IV with electrons (AA *vs*. Asc+TMPD, *p* < 0.001).

While mitochondrial respiration rates of platelet mitochondria were all significantly different from 0 (all *p* < 0.03; Fig. 2B), this was not the case for cell-free mitochondria (*OXPHOS*_*CI+II*_: *p* = 0.60; *LEAK* _*CI+II*_: *p* = 0.60; *ETS* _*CI+II*_: *p* = 0.25), except for complex IV respiration (*COX*: *p* = 0.028). Mitochondrial respiration rates of platelet mitochondria were significantly higher than cell-free ones (*OXPHOS*_*CI+II*_, *LEAK* _*CI+II*_ and *ETS* _*CI+II*_: all *p* < 0.03; Fig 2B), except for complex IV respiration (*i*.*e. COX*), for which cell-free mitochondria actually had a slightly higher respiration rate than platelet ones (*COX*: *p* = 0.046; Fig. 2B).

## Discussion

As noted by (7), the evidence provided by (4) was insufficient to infer the functionality or the respiratory competence of human circulating cell-free mitochondria. By conducting a comprehensive analysis of mitochondrial bioenergetics of human circulating cell-free mitochondria in direct comparison to mitochondria extracted from platelets of the same blood samples, we found no significant support for the functionality or the respiratory competence of those cell-free mitochondria. Indeed, there was no significant evidence that cell-free mitochondria possess a functional electron transport system since they exhibited no responses to mitochondrial substrates, ADP or inhibitors. Yet, when fueling directly complex IV with exogenous electrons (*i*.*e*. Asc+TMPD), a significant mitochondrial O_2_ consumption was detected (similarly to (4), except that no negative control was provided in their study). This shows that cell-free mitochondria retain a small fraction of the ETS, but that they are unlikely to be really functional or respiratory competent *in vivo* since electrons have to enter the ETS through complex I and/or II. In contrast, mitochondria isolated from platelets show the typical response to the SUIT protocol. Mitochondrial quantity per assay is unlikely to be a bias in the data being presented here since cell-free mitochondria only exhibited a slightly lower MitoTracker Green signal compared to mitochondria isolated from platelets, and even had a slightly higher complex IV activity (sometimes used as a proxy of mitochondrial quantity (12)). The absence of narrow-sense functionality (*i*.*e*. capability of oxidative phosphorylation) of circulating cell-free mitochondria suggested by our results is a key point regarding the identification of their potential physiological role(s). For instance, (8) hypothesized that extracellular mitochondria could help in restoring homeostasis by accumulating at sites of energy deficits, which is unlikely to be true if those mitochondria have no or an extremely low capacity for oxidative phosphorylation.

Yet, using flow-cytometry fluorescence assays, (6) recently reported that at least part of the cell-free mitochondria of human and murine blood could remain functional. Indeed, part of the cell-free mitochondria identified by flow cytometry and MitoTracker Green fluorescence appear capable of maintaining a mitochondrial membrane potential since they retained TMRE probe, and such retention was lowered by the addition of the mitochondrial uncoupler FCCP (6). Considering that the isolation protocol for cell-free mitochondria is likely to result in many microparticles not containing ‘intact’ mitochondria (6), one hypothesis could be that mitochondrial quantity is too low to enable the detection of significant functionality by high-resolution respirometry. Yet, considering the estimated density of functional cell-free mitochondria provided by (6) (*i*.*e*. 1.4 × 10^6^/mL of blood), the samples used in the present study should have contained *ca*. 110 × 10^6^ ‘intact’ extracellular mitochondria. Considering that human platelets contain on average 4.9 mitochondria per cell (13), mitochondrial respiration of our cell-free mitochondria samples should be approximately similar to the respiration of 22 × 10^6^ platelets. Mitochondrial respiration of human platelets has been reported by others to be around 0.35 pmol.s^-1^.10^6^ cells^-1^ (*OXPHOS*_*CI+II*_ in (14)). Thereby, if cell-free mitochondria from human blood had respiration rates in the same range as platelet mitochondria, we would expect our cell-free mitochondria to exhibit a respiration of *ca*. 7.6 pmol.s^-1^.mL^-1^, which is approximately 8 times above the detection limit of our high-resolution respirometry assay (*e*.*g. LEAK*_*CI+II*_ of platelet mitochondria was 7.3 ± 2.4 pmol.s^-1^.mL^-1^ in our study).

Although fully rejecting the hypothesis that human circulating cell-free mitochondria are functional would require a way larger sample size (*e*.*g*. N > 90 for reaching a statistical power of 0.80 for *OXPHOS*_*CI+II*_ based on data presented in Fig. 1B), our results do not support the functionality of circulating cell-free mitochondria. Further studies investigating the functionality or lack of functionality of circulating cell-free mitochondria are now required, since this appears as a critical point to elucidate their potential physiological role(s) (*e*.*g*. (8)).

## Acknowledgements

AS is grateful to Dr. T Hentinen and Prof. H Minn for helping to organize the study, to R Sjöroos for collecting blood, to the six volunteers for giving their blood, as well as to the ‘Turku Collegium for Science and Medicine’ for funding his research, and to the CafR team of La Piclière La Clusaz for hosting him while writing the initial commentary on (4).

## Author contribution

AS designed the study and conducted all the laboratory work, statistical analysis and writing of the manuscript.

